# Dysregulated gene expression of imprinted and X-linked genes: a link to poor development of bovine haploid androgenetic embryos

**DOI:** 10.1101/2020.10.23.350405

**Authors:** Luis Aguila, Jacinthe Therrien, Joao Suzuki, Mónica García, Amanda Trindade, Lawrence C. Smith

**Affiliations:** Département be biomedicine vétérinaire, Centre de recherche en reproduction et fertilité, Université de Montreal, Saint-Hyacinthe, QC, J2S 2M2, Canada

**Keywords:** uniparental embryo, paternally imprinting, epigenetic, bovine ICSI, diploid embryo, oocyte enucleation

## Abstract

Mammalian uniparental embryos are efficient models for genome imprinting research and allow studies on the contribution of the paternal and maternal genome to early embryonic development. In this study, we analyzed different methodologies for production of bovine haploid androgenetic embryos (hAE) to elucidate the causes behind their poor developmental potential. The results showed that hAE can be efficiently generated by using intracytoplasmic sperm injection and oocyte enucleation at telophase II. Although haploidy does not disturb early development up to around the 3^rd^ mitotic division, androgenetic development is disturbed after the time of zygote genome activation those that reach the morula stage are less capable to become a blastocyst. Analysis of gene expression indicated abnormal levels of methyltransferase 3B and key long non-coding RNAs involved in X-chromosome inactivation and genomic imprinting of the KCNQ1 locus, which is associated to the methylation status of imprinted control regions of XIST and KCNQ1OT1. Thus, our results seem to exclude micromanipulation consequences and chromosomal abnormalities as major factors in developmental restriction, suggesting that their early developmental constraint is regulated at an epigenetic level.

## 1. Introduction

In contrast to lower animal classes that can develop from a single parent by parthenogenesis, mammals have developed parental-specific epigenetic strategies such as genomic imprinting that require the contribution from the paternal and maternal genomes to develop fully to term. Nonetheless, early development can be achieved very efficiently from uniparental embryos in mammals using different artificial oocyte activation and/or micromanipulation techniques, which has been extremely useful in delineating genomic function, imprinting status and its role in ontogenesis (Cruz et al., 2008; Hu et al., 2015b).

Diploid androgenetic and gynogenetic/parthenogenetic embryos possess two sets of paternal or maternal genomes, respectively, while their haploid counterparts contain only one paternal or maternal genome. Although haploid development is a normal part of the life cycle for some animals (e.g., parasitic wasps), haploidy in mammals is restricted to gametes, which are structurally specialized for fertilization and mitotically incompetent (Shuai and Zhou, 2014).

Uniparental haploid embryos are efficient models for genome imprinting research and enable studies on the contribution of the paternal and maternal genome to early embryonic development. Moreover, haploid embryos have been used to derive embryonic stem cells and hold great promise for functional genetic studies and animal biotechnology (Panneerdoss et al., 2012; Kokubu and Takeda, 2014; Bai et al., 2016; Bai et al., 2019).

Although haploid embryonic stem cells have been obtained in several mammals (Leeb and Wutz, 2011; Yang et al., 2013; Zhong et al., 2016), most reports have indicated poor rates of blastocyst formation, suggesting impairments at early stages of embryonic development. In mice, studies have revealed that the preimplantation developmental potential of haploids is significantly impaired relative to diploid embryos due mainly to the disruption of gene regulatory mechanisms (Latham et al., 2002) and abnormal imprinted gene expression (Hu et al., 2015b). However, there are only a few studies characterizing the causes of limited development of haploid androgenetic embryos (hAE) in other mammals models, particularly in domestic species where the androgenetic embryonic stem cells would provide an useful route for genetic manipulations (Lagutina et al., 2004; Matsukawa et al., 2007; Park et al., 2009; Vichera et al., 2011).

The generation of mammalian hAE have been achieved by using a variety of methods. In mouse species the bisection of zygotes after fertilization (Tarkowski and Rossant, 1976), the removal of the maternal pronucleus from fertilized eggs at the pronuclear stage (Yang et al., 2012), and the injection of sperm into enucleated oocytes (Li et al., 2012; Yang et al., 2012) have been applied successfully. However, in bovine species the efforts to visualize and enucleate zygotes at pronuclear stages is hampered by the presence of dense lipid vesicles, and thus, usually, removal of the oocytes metaphase spindle is performed pre-IVF, as in the case of SCNT, at approximately 18 to 20 h after beginning of *in vitro* maturation (MII enucleation). Otherwise, during the telophase to anaphase transition of meiosis II the second polar body is a reliable indicator of the position of the oocyte’s spindle and it can be reliably used to enucleate mammalian oocytes (Bordignon and Smith, 1998; Kuznyetsov et al., 2007; Sagi et al., 2019).

Therefore, our aims were to establish an efficient method to produce bovine hAE and identify the potential causes of their severely limited developmental potential. Our results indicate that the developmental restriction of the androgenetic haploid embryos occurs at the time of the major transcriptional activation and it is associated with the altered expression of key epigenetically regulated genes. The significance and possible explanations for these findings are discussed.

## 2. Material and methods

### Oocyte collection and in vitro maturation

Bovine ovaries were obtained from a local slaughterhouse and transported to the laboratory in sterile 0.9% NaCl at 25–30°C in a thermos bottle. Cumulus–oocyte complexes (COCs) were aspired from 5 mm to 10 mm antral follicles using a 12-gauge disposable needle. For in vitro maturation (IVM), COCs with several cumulus cell layers were selected, washed and placed in maturation medium composed of TCM199 (Invitrogen Life Technologies), 10% fetal bovine serum (FBS), 0.2 mM pyruvate, 50 mg/mL gentamicin, 6 μg/mL luteinizing hormone (Sioux Biochemical), 6 μg/mL follicle-stimulating hormone (Bioniche Life Science) and 1 μg/mL estradiol (Sigma). In vitro oocyte maturation was performed for 22-24 h at 38.5°C in a humidified atmosphere at 5% CO_2_.

### Sperm preparation

Straws of non-sexed and sex-sorted semen stored in liquid nitrogen were thawed for 1 min in a water bath at 35.8°C, added to a discontinuous silane-coated silica gradient (45 over 90% BoviPure, Nidacon Laboratories AB), and centrifuged at 600 X g for 5 min. The supernatant containing the cryoprotectant and dead spermatozoa were discarded, and the pellet with viable spermatozoa was re-suspended in 1 mL of modified Tyrode’s lactate (TL) medium and centrifuged at 300 X g for 2 min.

### In vitro fertilization

After 222-24 h of IVM, COCs were washed twice in TL medium before being transferred in groups of 5 to 48 μl droplets under mineral oil. The in vitro fertilization (IVF) droplets consisted of modified TL medium supplemented with fatty-acid-free BSA (0.6% w/v), pyruvic acid (0.2 mM), heparin (2 μg/mL) and gentamycin (50 mg/mL). COCs were transferred to IVF droplets 15 min prior to adding the spermatozoa. To stimulate sperm motility, penicillamine, hypotaurine and epinephrine (2 mM, 1 mM and 250 mM, respectively) were added to each droplet. The selected spermatozoa were counted using a hemocytometer and diluted with IVF medium to obtain a final concentration of 1 × 10^6^ sperm/mL. Finally, 2 μL of the sperm suspension was added to the droplets containing the matured COCs. The fertilization medium was incubated at 38.5°C for 18 h in a humidified atmosphere of 95% air and 5% CO_2_. Presumptive zygotes were denuded by treatment with 0.1% bovine testicular hyaluronidase.

### Intracytoplasmic sperm injection

Intracytoplasmic sperm injections (ICSI) was performed according to standard protocols (Horiuchi et al., 2002) on the stage of a Nikon Ti-S inverted microscope (Nikon Canada Inc., Mississauga, ON, Canada) fitted with Narishige micromanipulators (Narishige International, Japan) and Piezo PMM 150HJ/FU (Prime tech Ltd., Japan). Before ICSI, oocytes were denuded of granulosa cells by gently pipetting in the presence of 1 mg/mL hyaluronidase, selected for the presence of the first polar body and randomly allocated to experimental groups. After ICSI, oocytes were washed at least three times and cultured in modified synthetic oviduct fluid (mSOF) media as previously described by Landry et al. (2016).

### Production of haploid embryos

Bovine haploid androgenetic embryos (hAE) produced by IVF were enucleated by removing the oocyte’s chromosomes (enucleation) either before or after insemination. When enucleating before IVF, COCs were denuded at 24 h after IVM, the oocytes were then exposed 15 min to 5 μg/mL cytochalasin B and 10 μg/mL Hoechst 33342 and a small portion (±10%) of the cytoplasm surrounding the first polar body was removed by aspiration into a micropipette. The aspirated cytoplasmic bleb was observed under UV light to ascertain whether the metaphase II spindle (MII) had been properly removed at enucleation. Oocytes in which enucleation was performed after IVF were removed from the fertilization droplet at different times after insemination, denuded of the cumulus cells by gentle pipetting and those presumptive zygotes with recently extruded second polar bodies were placed in cytochalasin B and Hoechst 33342 for 15 minutes as described above. A cytoplasm portion (±10%) surrounding the second polar body was aspirated from the oocyte, checked for the presence of a telophase-stage (TII) spindle, washed and returned to in vitro culture medium droplets.

On the other hand, bAhE produced by ICSI were obtain by removing the oocyte TII spindle after 4h post-ICSI. Enucleated zygotes were cultured as described above. Parthenogenetic embryos were produced according (Ock et al., 2003). Briefly, chemical oocyte activation was performed between 20 to 24 h after IVM by 5 min exposure to 5 μM ionomicyn (Calbiochem, San Diego, CA, USA). To obtain haploid parthenotes, ionomycin treatment was followed by incubation in 10 mg/mL cycloheximide (CHX) for 5 h, which enables complete extrusion of the second polar body. For diploid parthenogenotes, ionomycin-activated oocytes were exposed for 5 h to CHX and 5 mg/mL of cytochalasin B to inhibits the extrusion of the second polar body and, thereby, induce diploidization. After parthenogenetic activation, haploid or diploid parthenotes were washed and allocated to in vitro culture drops.

### In vitro culture

For in vitro culture, groups of 10 embryos were placed in droplets (10 μl) of modified synthetic oviduct fluid (mSOF) with non-essential amino acids, 3 mM EDTA, and 0.4% fatty-acid-free BSA (Sigma-Aldrich) under embryo-tested mineral oil. The embryo culture dishes were incubated at 38.5°C with 6.5% CO_2_, 5% O_2_, and 88.5% N_2_ in saturation humidity. Cleavage rate was recorded at 48 h (Day 2) of culture (IVF and ICSI = Day 0). Morula and blastocyst development rate were recorded on days 6 and 7 post-fertilization, respectively. Some haploid embryos were cultured for an extra 24 h to determine blastocyst rates at day 8 (192 h after ICSI). After assessment of development, embryos were either fixed for cell number evaluation or snap-frozen in liquid N_2_ and stored at −80 °C for RNA extraction.

### Assessments of pronuclear formation and total cell number

Pronuclear formation was assessed at 18-20 h after activation or fertilization and embryo quality was assessed on the basis of morphology and total cell number. Briefly, embryos at day 7 were classified morphologically as morula (compacted and >32 cells), early blastocyst (<200 μm), expanded blastocyst (>200 μm), and hatched blastocyst (after complete extrusion from the zona pellucida). Embryos at different stages were fixed overnight in paraformaldehyde and stained with Hoechst 33342 (10 μg/mL) for 15 min, and total number of cells and pronuclear formation were observed and analyzed by fluorescence microscopy (Axio Imager M1, Zeiss, Canada).

### Karyotype analysis

After culture in the in presence of 0.05 μg/mL of Colcemid (KaryoMax^®^Life Technologies, Carlsbad, CA, USA) for 5 h. Embryos were exposed to a hypotonic (0.75 M KCl) solution for 10 min to induce swelling. Subsequently, embryos were placed on a clean glass slide in a small volume of medium. Methanol–acetic acid solution (1:1; v/v) was dropped on the embryos while gently blowing with the slides placed under the stereoscope and allowed to dry for 15 min at room temperature. After drying, slides were stained with Hoechst 33342 (10 μg/mL) for 15 min. Chromosome spreads were evaluated at ×1000 magnification using oil immersion optics and fluorescence microscopy (Axio Imager M1, Zeiss, Canada). Embryonic cells were classified as haploid (n=30), diploid (n=60), or aneuploid (n≠ 30 or 60) according to the number of chromosomes.

### Gene specific bisulfite sequencing

Genomic DNA extraction and bisulfite treatment were done using a kit (EZDNA methylation-direct kit, Zymo research). Primers specific for bisulfite-converted DNA were designed within the DMR region of XIST (gene ID:338325) and KCNQ1OT1 (gene ID:112444897). KCNQ1OT1: F: GGTTAGAGGAGTATTTTGAAGAGA, R: TCAACCCTCTCAACCAATAA, and for XIST: F: TTTTGTTGTAGGGATAATATGGTTGA, R: TCATCTAATTCCATCCTCCACTAACT. Each PCR reaction was performed in triplicate. The PCR reaction was carried out in a final volume of 50 uL containing 1–2 uL of bisulfite-treated DNA, 0.2 uM each primer, 0.3 mM mixed dNTP, 1X PCR buffer, 1.5 mM MgCl2 with 2U of Platimun Taq DNA Polymerase (Invitrogen). The reactions were performed using an initial 2-min step at 94 °C followed by 45 cycles of 30 sec at 94 °C, 30 sec at 53 °C, 1 min at 72 °C, and a final 5-min step at 72 °C. The PCR products were resolved in 1.2% agarose gels, followed by purification using the QIAquick Gel Extraction kit (Qiagen). Purified fragments were pooled and subcloned in pGEM-T Easy Vector (Promega). 16 clones for each sample were picked and sequenced. Validation of the imprinted status of each DMR was performed as previously described by Lafontaine et al. (2020), by assessing methylation of sperm DNA (expected methylation > 90% or <10%) and fibroblast cell DNA (40-60% expected methylation).

### RNA extraction and RT-PCR

For analysis of gene expression, embryos were pooled for each stage of development: 15 of 8-cell embryos, 5 morulas, 3 blastocyts and each group was done in triplicate. Total RNA from the pooled embryos was extracted using the Arcturus PicoPure RNA Isolation kit (Lifetechnologies) and reverse transcribed into cDNA using SuperScript Vilo (Invitrogen). Quantitative RT-PCR was performed using the RotorGene SyBr Green PCR kit (Qiagen) in a Rotorgene Q PCR cycler under the following amplification conditions: 95°C for 5 min, followed by 40 cycles at 95°C for 5 secs and at 60°C for 10 secs. Primers were designed using Oligo6 software and the geometric means of three housekeeping genes (GAPDH, ACTB and SF3A) was used for normalization. The stability of the housekeeping genes across our samples was confirmed using Bestkeeper (Pfaffl et al.,2002). A list of all primers used can be found in Supplemental Table 1.

### Statistical analysis

Quantitative data sets are presented3 as means and standard deviation (± S.D) and analyzed using one-way ANOVA. Post hoc analysis to identify differences between groups was performed using Tukey test. Binomial data sets, such as pronuclear formation, were analyzed by using Fisher test. Differences were considered significant at p < 0.05.

## 3. Results

### Haploid androgenetic embryos produced by enucleation after fertilization leads to better development, but it is unreliable due to high polyspermy levels

Since the resumption of meiosis in the oocyte by the fertilizing spermatozoa can vary significantly when using conventional IVF, oocytes were exposed to spermatozoa during different time periods to identify an optimal fertilization time point at which the second polarbody could be used to locate the spindle for enucleation. With this purpose, we evaluated developmental potential after removing presumptive zygotes from the IVF drops at different times after insemination. Results indicated that second polar bodies were present in 80% of the oocytes by 6 h or more post insemination (hpi) while only 50% were fertilized with insemination periods of 4 hpi or less. Moreover, removal the presumptive zygotes from the IVF drop after 6 hpi led to better preimplantation development when compared to the shorter exposure periods to spermatozoa (*p* < 0.05), indicating that a 6 hpi period was suitable for oocyte enucleation post fertilization (data not shown).

Having identified an optimal period to expose oocytes to spermatozoa for enucleations during extrusion of the second polar body, we performed an experiment to compare the developmental outcome of putative haploid zygotes that were denuded and enucleated either pre- ot post-IVF. Diploid controls, i.e., denuded but non-enucleated pre- and post-fertilization groups, were cultured concomitantly. Confirming previous results (Lagutina et al., 2004; Vichera et al., 2011), cleavage rates at 48 h did not differ between putative haploid and control diploid embryos, indicating that the first cell divisions are not affected by haploidy or the timing of enucleation. However, blastocyst development was significantly reduced in haploid embryos, indicating that adrogenetic haploidy disturbs early development beyond the first cleavage. Nonetheless, we found that instead of performing enucleation at MII, enucleations after IVF produced significantly more 8-cell (*p* < 0.05) at Day 2 and blastocyst (*p* < 0.01) stage embryos at Day 7 after IVF, indicating that enucleation of the oocyte’s spindle before fertilization is more detrimental to the development of haploid androgenetic embryos (Table 1).

**Table 1.**
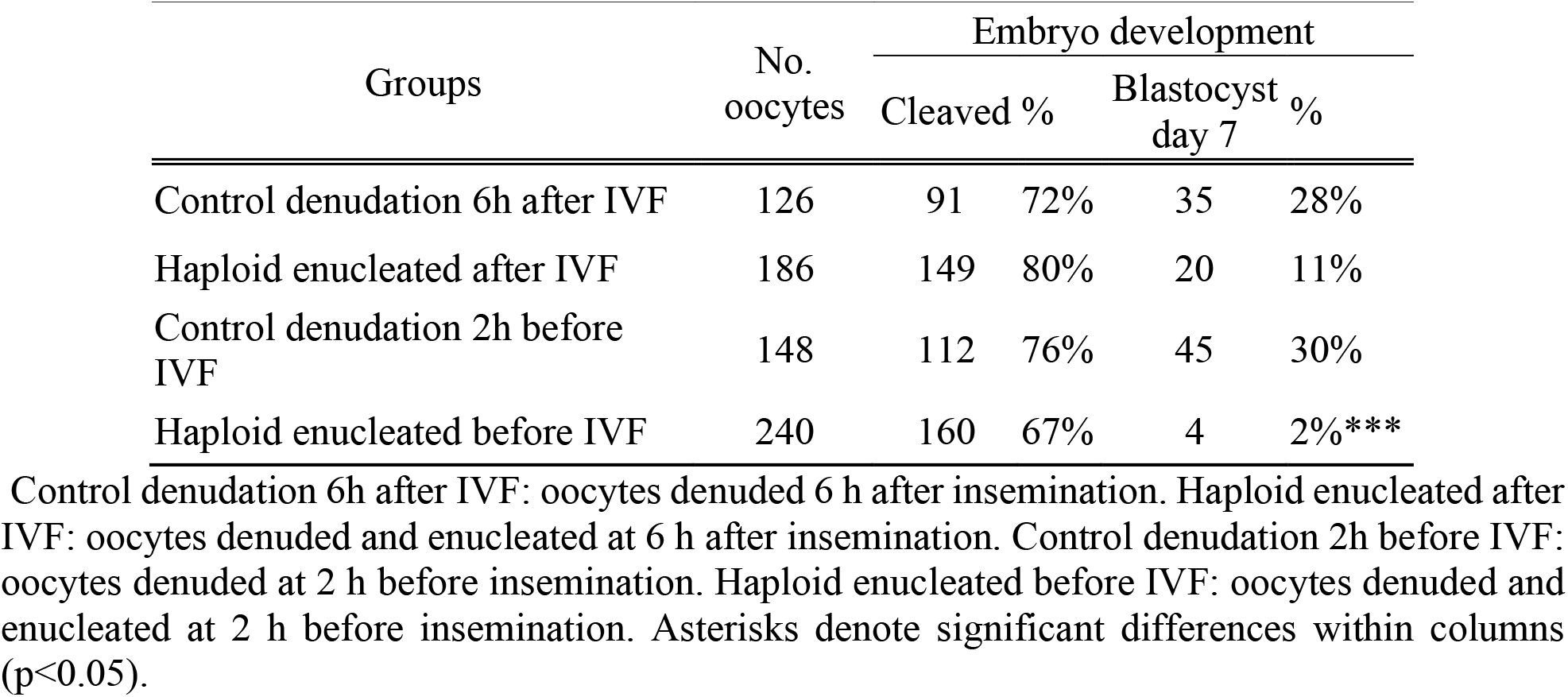
Development to cleavage and blastocyst stages at Day 2 (48 h) and Day 7 (168 h) post insemnination (hpi) of control and putative androgenetic embryos manipulated both before (pre-IVF) and after (post-IVF) in vitro fertilization (IVF).

| Groups | No. oocytes | Embryo development |  |  |  |
| --- | --- | --- | --- | --- | --- |
|  |  | Cleaved % |  | Blastocyst day 7 % |  |
| Control denudation 6h after IVF | 126 | 91 | 72% | 35 | 28% |
| Haploid enucleated after IVF | 186 | 149 | 80% | 20 | 11% |
| Control denudation 2h before IVF | 148 | 112 | 76% | 45 | 30% |
| Haploid enucleated before IVF | 240 | 160 | 67% | 4 | 2%*** |

Control denudation 6h after IVF: oocytes denuded 6 h after insemination. Haploid enucleated after IVF: oocytes denuded and enucleated at 6 h after insemination. Control denudation 2h before IVF: oocytes denuded at 2 h before insemination. Haploid enucleated before IVF: oocytes denuded and enucleated at 2 h before insemination. Asterisks denote significant differences within columns ( $p < 0.05$ ).

Next, we performed DNA staining of the putative haploid zygotes at 20 h after insemination to examine the number of procuclei of control and enucleated groups (Table 2). Since the presence of more than one pronucleus in enucleated and more than two pronuclei in control zygotes is indicative of polyspermy and/or mitotic errors which causes uncertainty with regard to ploidy in presumptive haploid (parthenogenetic and androgenetic) zygotes. No significant differences were observed in the level of multinucleated zygotes, neither between oocytes enucleated before and after-IVF (17% vs. 34%, *p* = 0.08), nor between enucleated and control groups (*p* > 0.07). Nonetheless, since appoximately one fifth and one third of the putative haploid zygotes derived from encucleated oocytes before and after-IVF, respectively, contained two or more prunuclei. Therefore, since these results indicated clearly that the production of bovine hAE by conventional IVF protocols leads to significant uncertainty with regard to ploidy, an unmistakable method was required to efficiently eliminate the possibility of polyspermic fertilization when deriving haploid androgenetic zygotes.

**Table 2.** Formation of pronuclei of zygotes fixed at 20 h after insemination in control and enucleated oocytes manipulated either before or after insemination.

| Group | Oocytes<br>(n) | Pronuclear Formation 20 hpi (No. and %) |  |  |  |  |  |  |
| --- | --- | --- | --- | --- | --- | --- | --- | --- |
|  |  | 1 PB |  |  | Others |  |  | *Multinucleated |
|  |  | 1 PN | 2 PN | PDSH | CSH | ≥ 3 PN | 0 PN |  |
| Control denuded 6h after IVF | 64 (3) | 0 (0%) | 44 (69%)* | 0 (0%) | 0 (0%) | 14 (22%) | 6 (9%) | 14 (22%) |
| Haploid enucleated after IVF | 80 (3) | 43 (54%)* | 6 (8%) | 0 (0%) | 0 (0%) | 21 (25%) | 10 (13%) | 27 (34%) |
| Control denuded 2h before | 10 (2) | 0 (0%) | 8 (80%)* | 0 (0%) | 0 (0%) | 1 (10%) | 1 (10%) | 1 (10%) |
| Haploid enucleated before IVF | 12 (2) | 9 (75%)* | 0 (0%) | 0 (0%) | 0 (0%) | 2 (17%) | 1 (8%) | 2 (17%) |

PB: polar body; PN = pronuclei; PDSH = partially decondensed sperm head; CSH = condensed sperm head. Control denudation 6h after IVF: oocytes denuded 6 h after insemination. Haploid enucleated after IVF: oocytes denuded and enucleated at 6 h after insemination. Control denudation 2h before IVF: oocytes denuded at 2 h before insemination. Haploid enucleated before IVF: oocytes denuded and enucleated at 2 h before insemination. \*Multinucleated: multinucleated zygote was assumed as the presence of more than one pronucleus in enucleated and more than 2 pronuclei in control zygotes. Asterisks denote significant differences within columns (p<0.05).

### Haploid androgenetic embryos can be obtained reliably by intracytoplasmic sperm injection and enucleation of the telophase II spindle

Due to the unreliability of conventional IVF in deriving x, we next examined the use of intracytoplasmic sperm injection (ICSI) toeliminate the posibility of polyspermic fertilization. Since better development was achieved by enucleation after IVF, we performed ICSI followed by enucleation 3-4 h later, i.e. when the telophase-II spindle and the second polar body were easily identified for microsurgical removal. In order to verify the efficiency of the enucleation procedure after ICSI, we evaluated the rate of pronuclear formation at 20 h post ICSI (Figure 1; Table 3) including the parthenogenetic haploid zygotes as a positive control for prescence of only one pronucleus.

**Figure 1.**
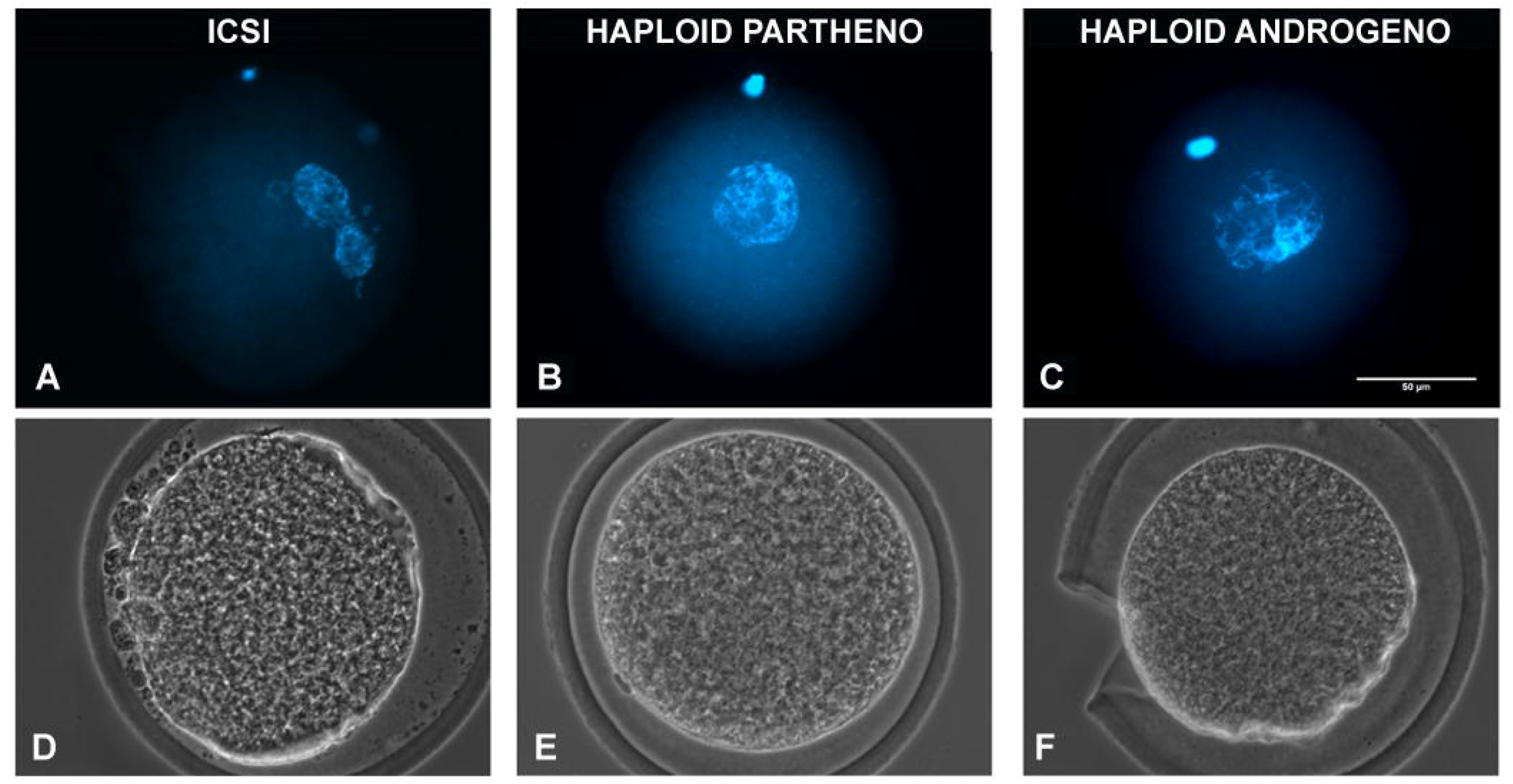
Representative images of a 1-cell stage zygote fixed at 20 h after activation showing DNA staining (upper) and phase-contrast images (lower) of a (a,b) ICSI, biparental embryo obtained by ICSI (2 pronuclei), (c,d) Haploid partheno, haploid parthenogenetic embryo (1 female pronucleus) obtained by oocyte activation using ionomycin followed by cyclohexymide, and (e,f) Haploid androgeno, haploid androgenetic embryo (1 male pronucleus) obtained by ICSI + oocyte enucleation. ICSI, intracytoplasmic sperm injection using female-sorted semen. Scale bar = 50 μm.

**Table 3.**
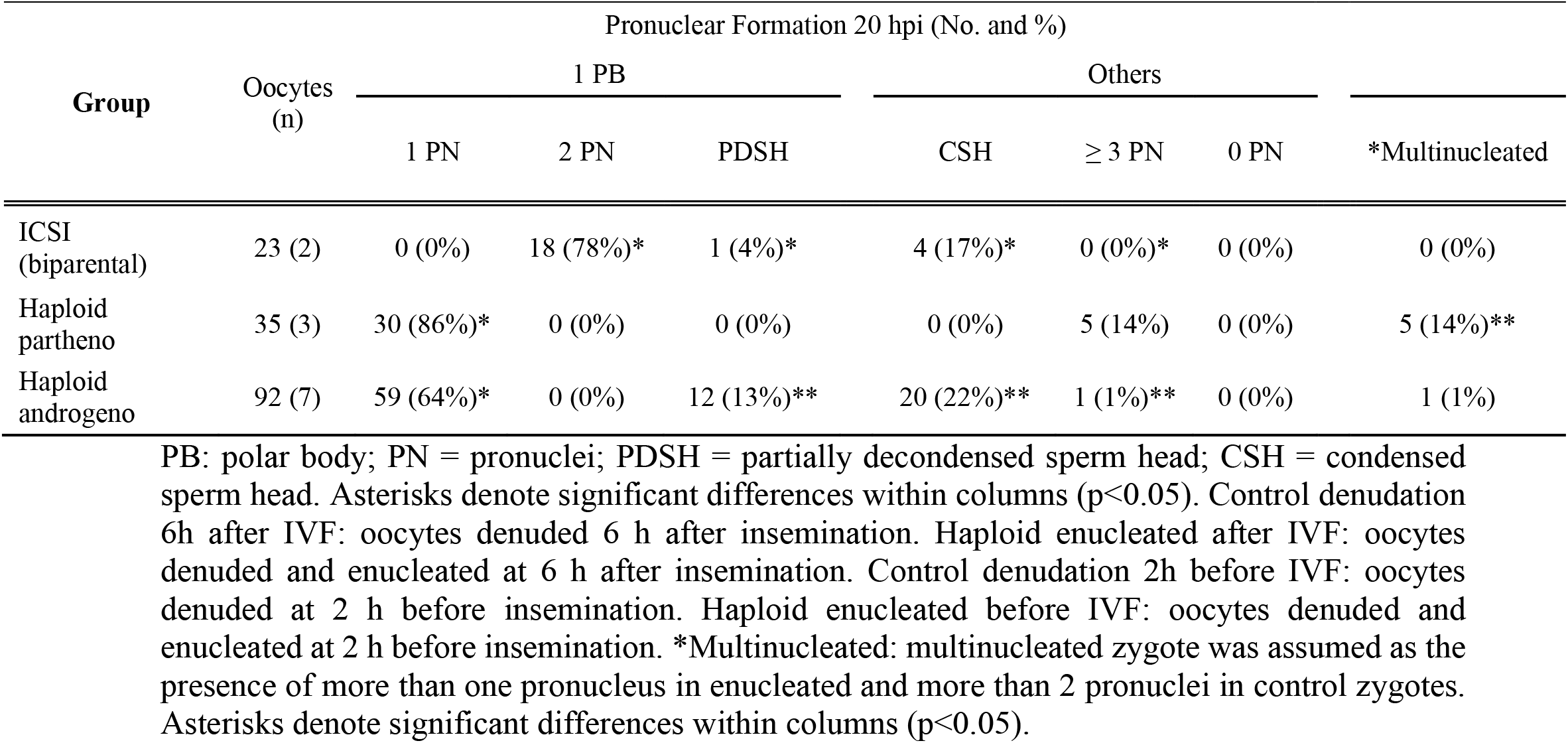
Formation of pronuclei and other chromatin structures in diploid (ICSI) and haploid zygotes (parthenogenetic and androgenetic) observed at 20 h after activation.

DNA staining showed that 99% of the enucleated zygotes after ICSI had only one chromatin structure (Table 3, Figure 1). Additionally, there was a tendency for enucleated oocytes to not support the complete decondensation of paternal chromatin and formation of a pronucleus when compared to ICSI controls (35% vs. 21%, respectively), suggesting that the removal of the telophase spindle at first hours after ICSI disturbs sperm-head decondensation. Together, these results confirm that the use of ICSI is a more reliable approach to derive bovine haploid androgenetic zygotes.

### Haploid androgenetic embryos develop poorly and slowly to the blastocyst stage

Once the reliability of the ICSI approach for deriving hAE was verified, we next compared early developmental rates of haploid and diploid control groups at different times of in vitro culture. Because previous reports have shown that androgenetic embryos produced using Y chromosome-carrying sperm are unable to support development to the blastocyst stage (Latham et al., 2000; Latham et al., 2002; Yang et al., 2012), we used semen that had been sorted (sexed) to obtain sperm with an X-chromosome or a Y-chromosome. Except for the IVF group that showed the highest cleavage rate (90%; p < 0.01), all the remaining groups showed similar levels of cleavage (range 72% to 74%) (Table 4). However, the rate of embryos having ~8 cells at 48 h of culture (suppl. Fig. 1) was lower only in haploid parthenogenetic embryos, suggesting that manipulation procedures involved in generating hAE do not affect early cleaving. Development up to morula and blastocyst was similar among the diploid IVF and ICSI controls. The haploid parthenogenetic group showed lower blastocyst rate than biparental embryos, but higher than the hAE (Table 4). On the other hand, hAE showed the lowest developmental potential (9% and 3% for morula and blastocyst stage, respectively; p<0.0001) compared to biparental and the haploid parthenogenetic group (Table 4), indicating that androgenetic haploidy is lees suitable for support preimplantation development compared to the parthenogenetic haploidy. Besides, only hAE produced with sperm carrying X-chromosome reached the morula and blastocyst stages (Table 4). DNA staining showed that hAE produced with sperm carrying Y-chromosome did not develop beyond 20 cells (suppl. Fig. 2). Further assessment of embryo morphology at Day-7 indicated major differences between androgenetic and the remaining groups (Figure 2). Moreover, assessment of nuclear number of Day-6 morulae and Day-7 blastocysts showed that androgenotes contained significantly fewer cells when compared to ICSI control and haploid parthenotes of the same age (Figure 3). Actually, some androgenetic embryos only reached the blastocyst at Day-8, indicating that the blastulation is delayed in the few hAE that are able to reach the blastocysts stage (Data not shown). After cleavage, hAE underwent developmental arrest concurrently with time of zygote genome activation, where only 13% of the cleaved embryos progressed up to morula stage, compared to haploid parthenotes (32%) and diploid groups (>40%) (Table 4). In addition, haploid androgenotes arrested once again at morula stage when only 26% of the Day-6 morula became a blastocyst (Table 4). Since the ratio of haploid parthenogenetic morulas (68%) that become blastocyst was significantly higher (p<0.005) than the haploid androgenetic group, these results indicate that, as in the other mammalian models, the bovine haploid paternal condition is less capable to support early embryonic development when compared to its maternal counterpart.

**Table 4.** Development to cleavage (Day-2) and blastocyst (Day-7) stages of embryos produced by in vitro fertilization (IVF), intracytoplasmic sperm injection (ICSI) using non-sexed and sexed sperm, haploid parthenogenetic and androgenesis using sexed spermatozoa.

| Group | Oocytes<br>(n) |  | Cleaved<br>embryos 48<br>hpi (%) |  | Embryo development |  |  |  |  |
| --- | --- | --- | --- | --- | --- | --- | --- | --- | --- |
|  |  |  |  |  | Morulas/<br>oocyte |  | Blastocyst/<br>oocyte |  | Blastocyst/<br>morula |
| IVF | 164 | (10) | 148 | 90%** | 59 | 36% | 48 | 29% | 81% |
| ICSI X-carrying | 260 | (16) | 187 | 72% | 80 | 31% | 68 | 26% | 85% |
| ICSI Y-carrying | 189 | (5) | 122 | 65% | 50 | 26% | 45 | 24% | 90% |
| Haploid parthenote | 354 | (16) | 261 | 74% | 84 | 24% | 57 | 16%* | 68%** |
| Haploid andro-X | 359 | (17) | 262 | 73% | 34 | 9%** | 9 | 3%**** | 26%**** |
| Haploid andro-Y | 146 | (5) | 103 | 71% | n.a | n.a | n.a | n.a | na |

ICSI: intracytoplasmic sperm injection using non-sexed sperm. ICSI X-carrying: intracytoplasmic sperm injection using X-chromosome sexed sperm. Haploid andro-X: haploid androgenetic embryos generated using X-chromosome sexed sperm. Haploid andro-Y: haploid androgenetic embryos generated using Y-chromosome sexed sperm n.a. data is not available. Asterisks denote significant differences within columns (\* $p < 0.05$ ; \*\* $p < 0.005$ , \*\*\*\* $p < 0.0001$ ).

**Figure 2.**
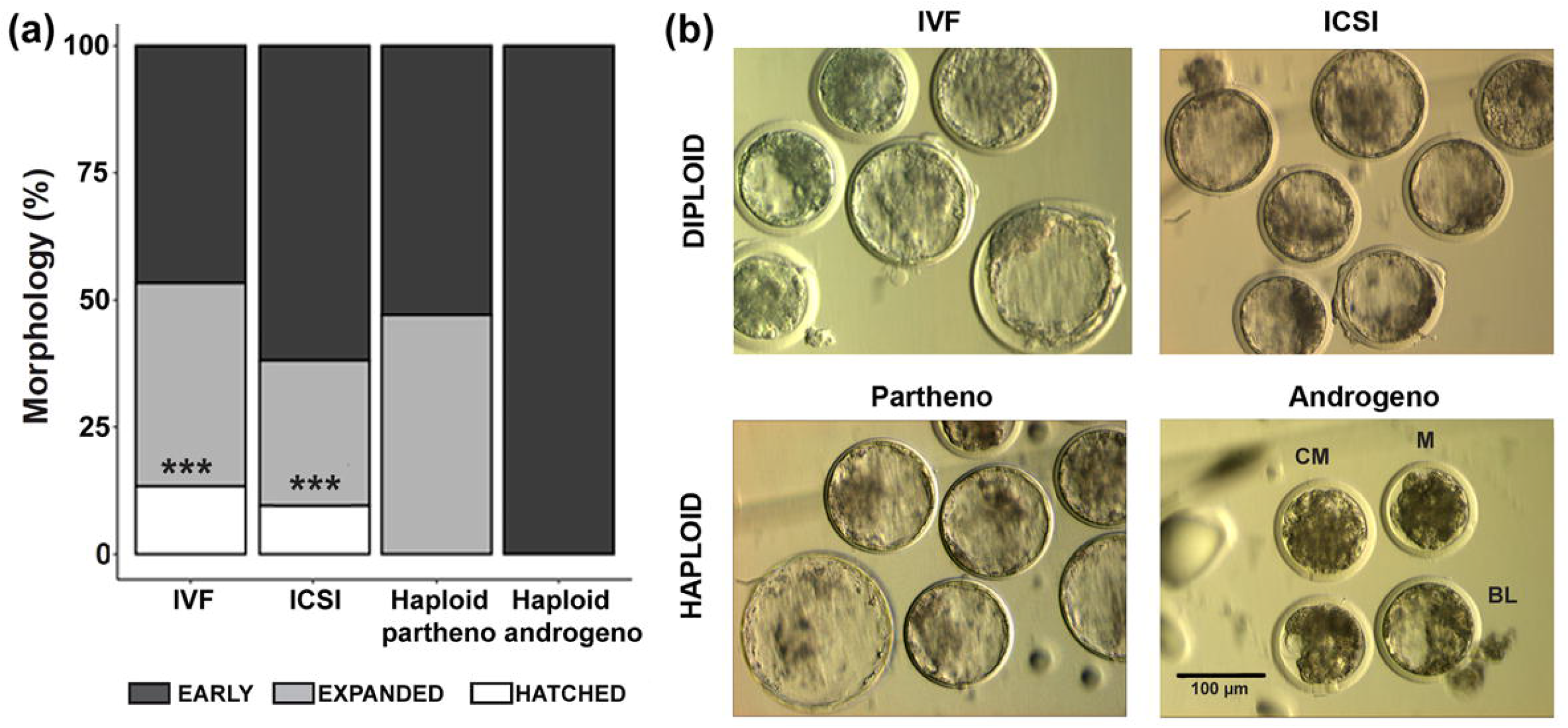
Morphological assessment of haploid and diploid embryos at Day-7 (168 h) of culture. (a) Percentage of different blastocyst stages. (b) Representative images of the most advanced embryos from different controls and haploid groups. IVF, in vitro fertilized; ICSI, intracytoplasmic sperm injection using female-sorted semen; Haploid partheno, haploid parthenogenetic embryos obtained by oocyte activation using ionomycin followed by cyclohexymide; Haploid androgeno, haploid androgenetic embryo obtained by ICSI + oocyte enucleation. M: compact morula; CB: cavitating blastocyst; BL: blastocyst. Scale bar = 100 μm.

**Figure 3.**
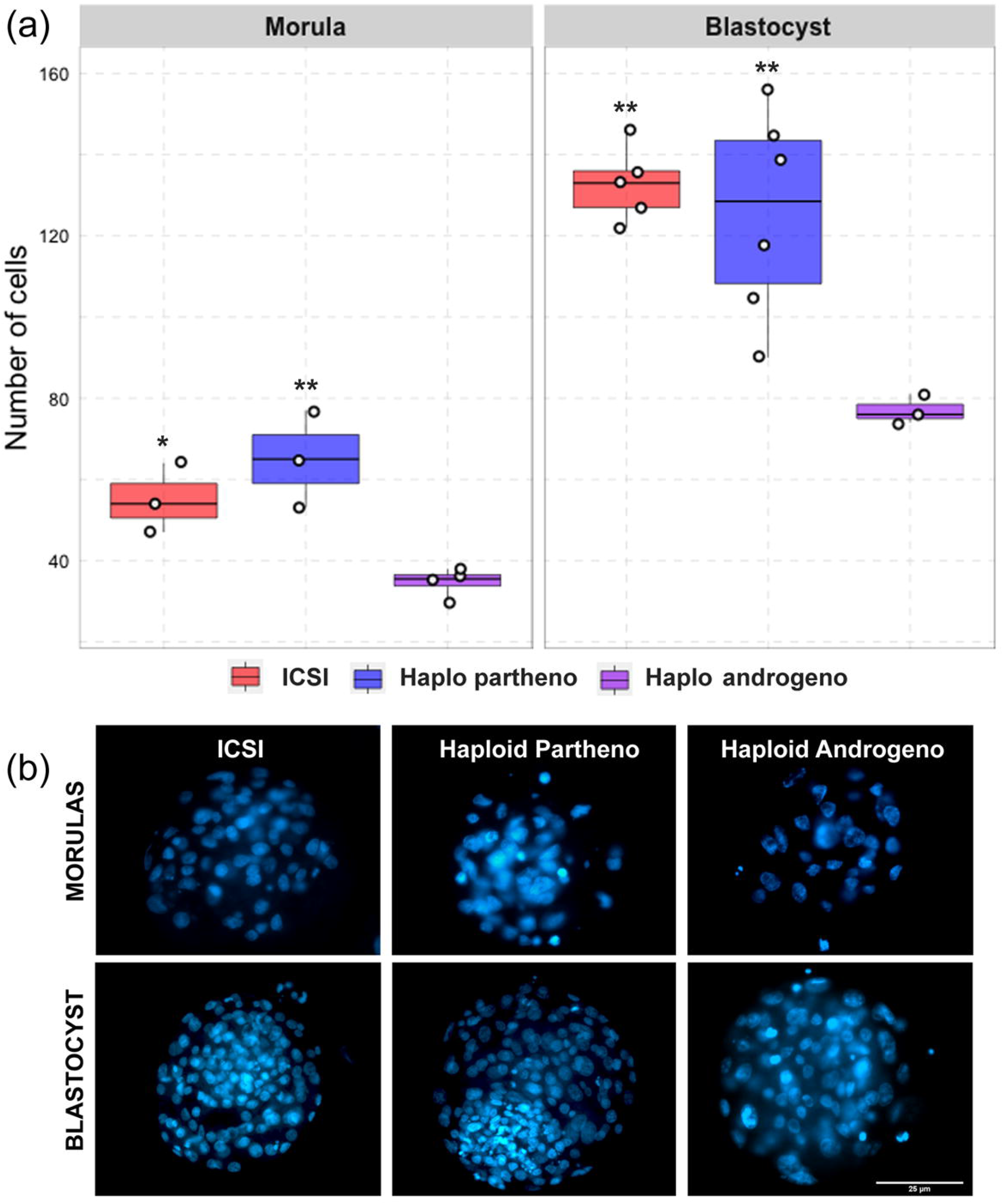
Cell number of diploid and haploid morula and blastocyst stage embryos. (a) Nuclear counts and (b) representative images of morula and blastocyst stage embryos harvested at Day-6 (144 h) and 168 h (day 7) of culture, respectively. IVF, in vitro fertilized; ICSI, intracytoplasmic sperm injection using female-sorted semen; Haploid partheno, haploid parthenogenetic embryos obtained by oocyte activation using ionomycin followed by cyclohexymide; Haploid androgeno, haploid androgenetic embryo obtained by ICSI + oocyte enucleation. Scale bar = 25 μm.

### Haploid androgenetic embryos maintain stable ploidy

Chromosomal anomalies have been identified in embryos handled in vitro (Kawarsky et al., 1996; Rubio et al., 2003; Ross et al., 2008). Therefore, we decided to verify whether the ploidy of the haploid and diploid embryos was particularly disturbed through karyotyping of metaphase-arrested cells. Surprisingly, most of the analyzed haploid androgenetic blastomeres (81%) contained normal haplotype (X=30), which contrasted, but not significatively (p > 0.05), with the diploid ICSI and haploid parthenogenetic embryos, that contained fewer (46% and 35%, respectively) normal karyotypes (Table 5; Figure 4).

**Table 5.** Chromosomal composition of bovine biparental diploid ICSI and haploid uniparental embryos

| Group | Embryos<br>evaluate<br>d (cells) | No. of cells |  |  |  |
| --- | --- | --- | --- | --- | --- |
|  |  | 1n | 2n | Aneuploid | Total<br>abnormal |
| ICSI | 10 (28) | 1 (4%)<sup>a</sup> | 13 (46 %) | 14 (50%) | 15 (54%) |
| Haploid<br>partheno | 20 (42) | 15 (35%)<sup>ab</sup> | 8 (19%) | 19 (45%) | 27 (64%) |
| Haploid<br>Androgeno | 9 (11) | 9 (81%)<sup>bc</sup> | 0 (0%) | 2 (19%) | 2 (19%) |

ICSI: intracytoplasmic sperm injection using X-sexed sperm. Androgenetic haplo-X: androgenetic haploid embryos generated using X-chromosome sexed sperm.

**Figure 4.**
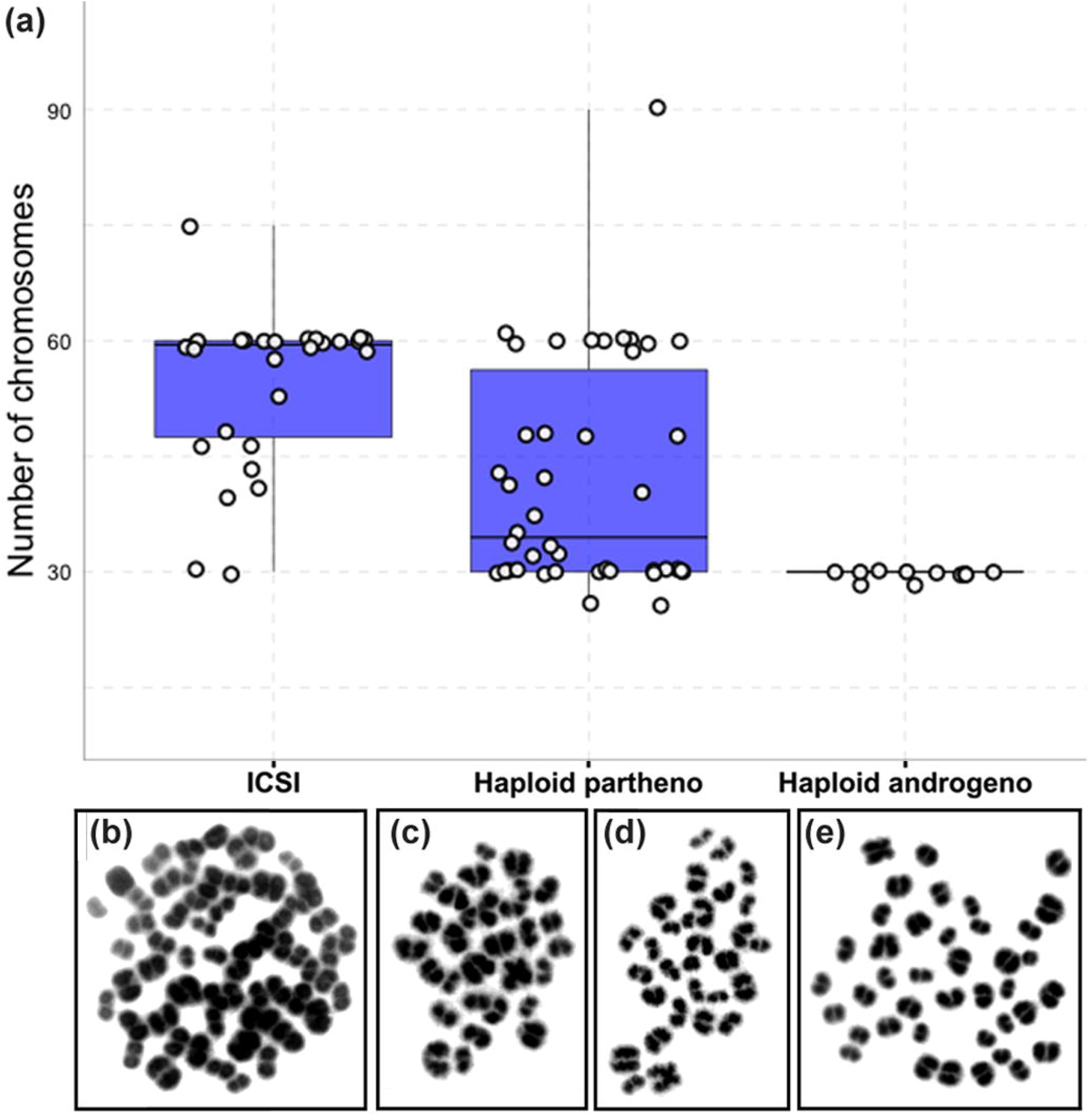
Karyotype analysis of haploid and diploid morula stage (Day-6) embryonic blastomeres. (a) Group chromosomal number distributions showing mean values (blue horizontal lines) with standard deviations (red horizontal lines). (b-e) Representative images of DAPI-stained chromosomal spreads of embryonic blastomeres from (b) diploid ICSI (60 chromosomes) (c) haploid parthenote (30 chromosomes), (d) haploid androgenote (30 chromosomes), and (e) aneuploid (40 chromosomes) parthenote embryo. ICSI, intracytoplasmic sperm injection using female-sorted semen; Haploid partheno, haploid parthenogenetic embryos obtained by oocyte activation using ionomycin followed by cyclohexymide; Haploid androgeno, haploid androgenetic embryo obtained by ICSI + oocyte enucleation

### Early cleavage events are not affected in hAE

In humans, developmental anomalies during the first mitotic divisions of in vitro-derived embryos have been associated with poor gamete qualities and in vitro processing (Hardy et al., 1993; Pelinck et al., 1998; Alikani et al., 2000; Babariya et al., 2017). In order to elucidate the anomalies associated with the poor development of haploid hAE, we first evaluated nuclear morphology of embryos that arrested between 1- to 3-cell stage after 48 h of culture. Apart from IVF-derived controls, all groups that underwent micromanipulation, such as ICSI, enucleation and chemical oocyte activation, had higher rate of mitotic anomalies (*p* < 0.001; Figure 5), suggesting that extensive in vitro manipulation of the oocyte is associated with early-stage developmental anomalies. Particularly, haploid androgenotes did not present additional anomalies when compared to ICSI-derived embryos, indicating that removal of the oocyte’s spindle at telophase does not further interfere with early cleavage. Together, these results indicate that mitotic errors during first cleavage divisions (i.e. 2^nd^ and 3^rd^) were caused mostly by the micromanipulation procedures, confirming that the inability to progress up to blastocyst by hAE arises mainly after the zygote genome activation.

**Figure 5.**
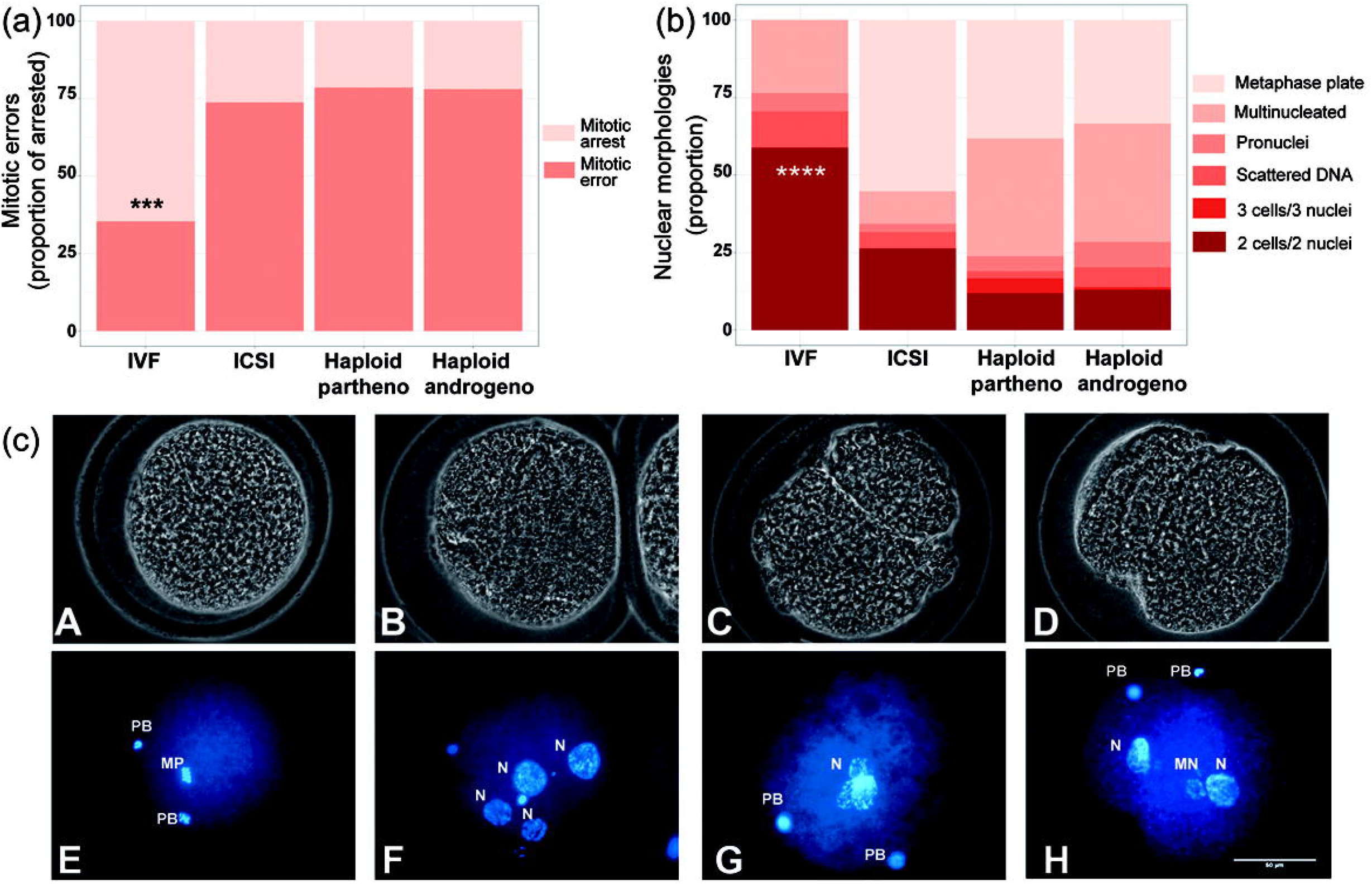
Developmental patterns of arrested embryos at 2- to 3-cell stage. (a) Proportion of embryos showing mitotic errors in relation to total number of arrested embryos. (b) Nuclear morphologies found in the zygotes showing developmental arrest. (c) Representative images of developmentally arrested embryos; (A, E) non-activated oocyte, (B, F) multinucleated zygote, (C, G) Anucleate blastomere, (D, H) Micronuclear formation. PB, polar body; mp, methaphase plate; n, nucleous; mn, micronucleous; ICSI, intracytoplasmic sperm injection using female-sorted semen; Haploid partheno, haploid parthenogenetic embryos obtained by oocyte activation using ionomycin followed by cyclohexymide; Haploid androgeno, haploid androgenetic embryo obtained by ICSI + oocyte enucleation. Scale bar 50 μm.

### Altered gene expression of X-linked genes and the KCNQ1 locus in hAE

Since haploid embryos possess exclusively maternal or paternal-derived chromosomes, genomic imprinting (autosomal or sex-related imprinting) offers another possible explanation to their poor early development (Latham et al., 2002). For instance, imprinting of the paternal X chromosome could potentially lead to development anomalies in haploid X chromosome-bearing androgenotes. Because of this, we analyzed the expression of X-linked genes and some genes previously described (Jiang et al., 2015) to undergo genomic imprinting in bovine species. To do this, we used two different stages, at at 8- to16-cell and morula stage embryos, basically to evaluate the expression levels at the time of the zygote genome activation (ZGA), and the most advanced stage of development available in haploid androgenotes (development to the blastocyst stage was seriously limited in this group). In addition, to analyze the effects of sex and ploidy on the gene expression levels, we included different control groups, such as ICSI female (ICSI using X-chromosome carrying sperm), ICSI male (ICSI using Y-chromosome carrying sperm), and parthenotes (both, diploid and haploid). At the time of ZGA (72 hpi), results indicate that the expression patterns of the X-linked genes XIST, PGK1 and HPRT were similar between haploid androgenotes, haploid parthenotes and biparental male and female embryos (Figure 6). In contrast, while the IGF2R imprinted gene did not show variations among groups, imprinted genes belonging to the KCNQ1 locus showed significant differences in expression. KCNQ1OT1, the paternally expressed long non-coding RNA involved in regulating the KCNQ1 locus, was significantly upregulated in haploid androgenotes compared to parthenotes (haploids and diploids) and biparental female embryos (Figure 6). Parthenogenetic embryos barely expressed KCNQ1OT1, confirming its imprinted nature for exclusive paternal expression. Similarly, CDK1 was upregulated in hAE. However, PHLDA2 showed lower levels in haploid androgenotes only when compared to diploid parthenotes (Figure 6).

**Figure 6.**
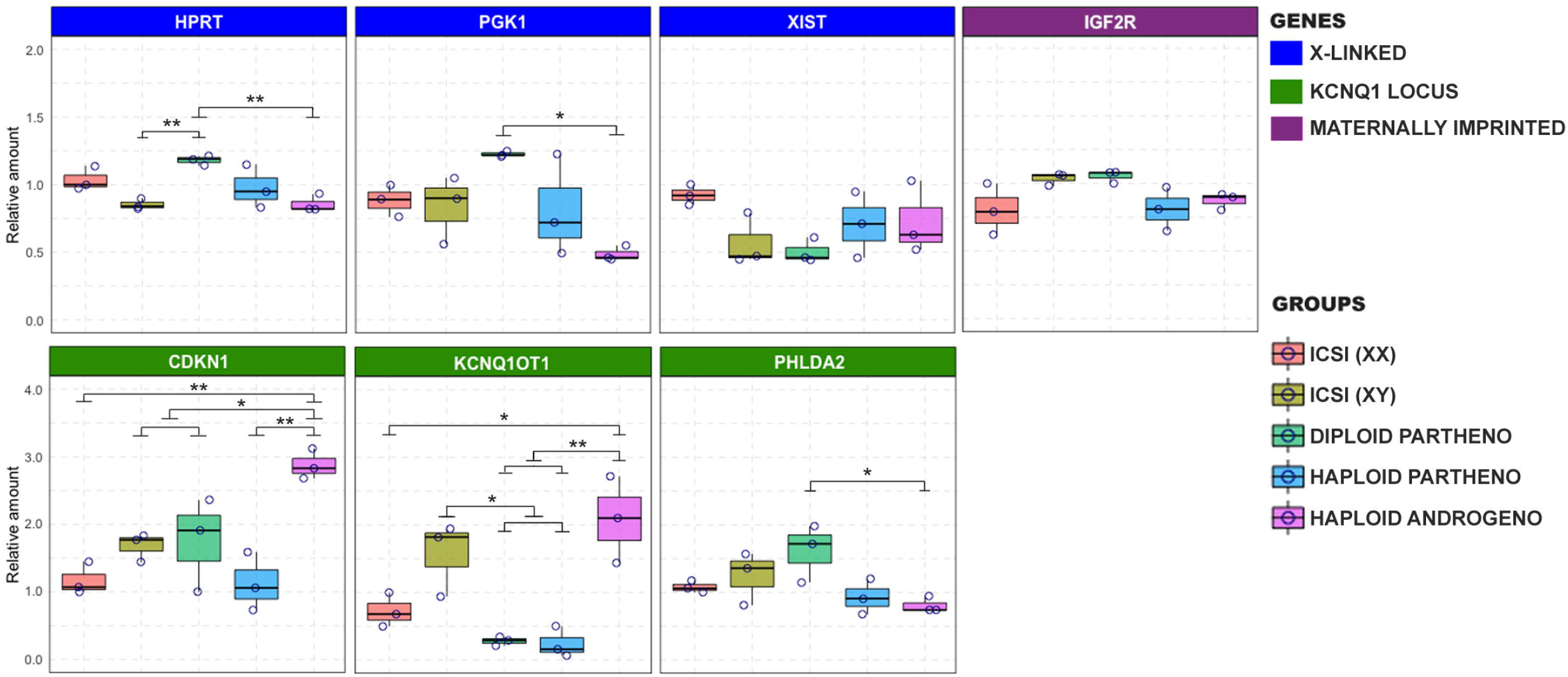
Relative gene expression of different imprinted and X-chromosome linked genes on haploid and diploid 8-cell stage embryos obtained by ICSI, parthenogenetic activation or ICSI + enucleation. Blue boxes: X-linked genes XIST, PGK1 and HPRT on Chromosome X; Green boxes: genes on the KNCQ1 locus KCNQ1OT1, CDKN1 and PHLDA2 (The three housekeeping genes used were GAPDH, ACTB and SF3A).

At the morula stage of development (day 6), effects on the expression of X-linked and the KCNQ1 imprinted locus were even further exacerbated. XIST transcript levels were significantly overexpressed in the haploid androgenotes compared to all the control groups, and PGK1 levels were higher haploid androgenotes compared to male biparental embryos (Figure 7). As for the KCNQ1 imprinted locus, haploid androgenotes showed significant upregulation of KCNQ1OT1 and PHLDA2 in comparison to parthenotes and biparental groups, whereas CDK1NC levels were unaffected (Figure 7). In contrast, expression patterns of the imprinted genes IGF2R and GNAS were not altered in haploid androgenotes, indicating that not all imprinted loci are disturbed in androgenotes. Altogether, the results show that hAE have altered gene expression of X-chromosome genes and imprinted genes from the KCNQ1 locus, suggesting that the developmental anomalies observed in haploid androgenotes at early stage of embryogenesis, i.e. at and soon after ZGA, are regulated at an epigenetic level.

**Figure 7.**
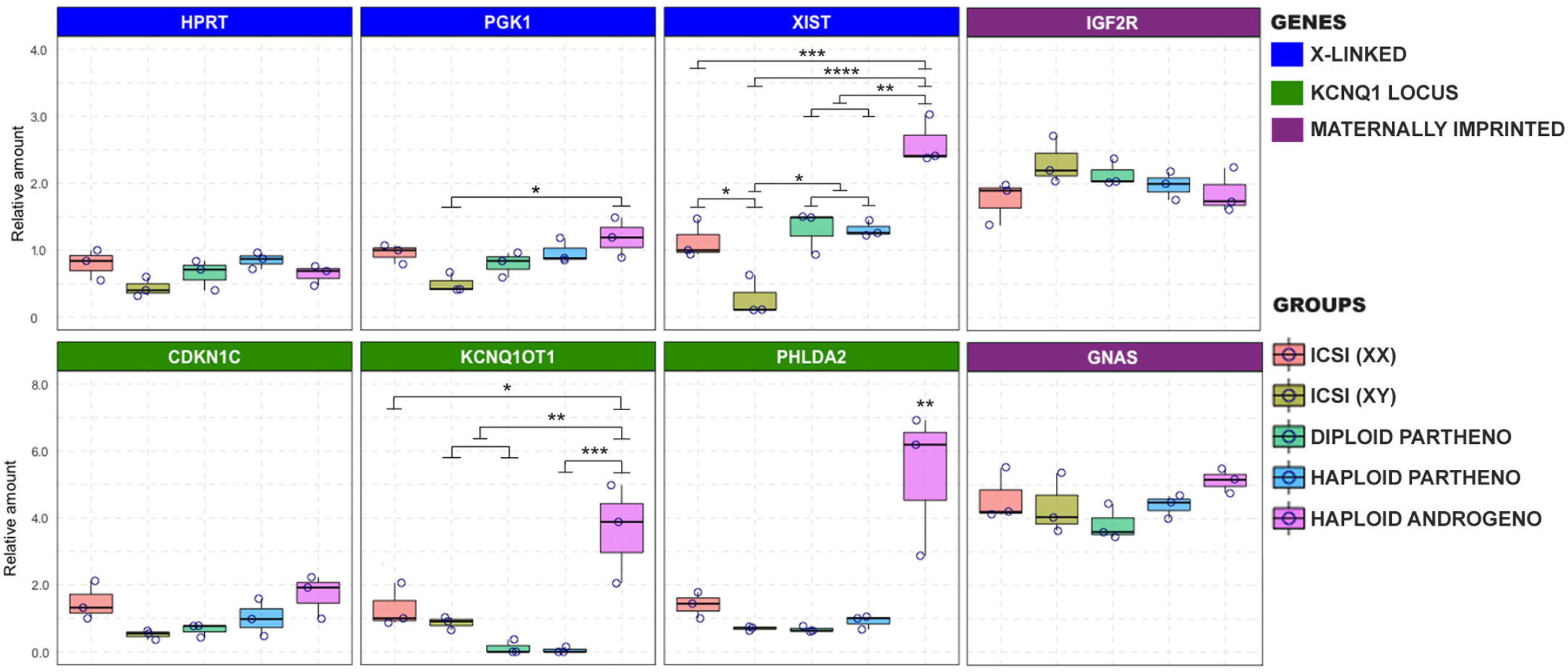
Relative gene expression of different imprinted and X-chromosome linked genes on haploid and diploid morula stage embryos obtained by ICSI, parthenogenetic activation or ICSI + enucleation. Blue boxes: X-linked genes XIST, PGK1 and HPRT on chromosome X; Green boxes: genes on the KNCQ1 locus KCNQ1OT1, CDKN1 and PHLDA2 (housekeeping genes used for normalization were GAPDH, ACTB and SF3A).

### Dnmt3b expression is downregulated in hAE

Genes expression is often regulated by DNA cytosine methylation, catalyzed by DNA methyltransferases, and it is often altered during in vitro culture (Lafontaine et al., 2020). Owing to the dysregulated transcript expression observed in hAE, specifically in genes from the KCNQ1 locus and X chromosome, we analyzed the expression of enzymes related to maintenance (DNMT1) and de novo (DNMT3B) DNA methylation as well as demethylation (TET1) in haploid (parthenogenetic and androgenetic) and biparental (ICSI) morula stage embryos. Although no differences were observed between the levels of DNMT1 and TET1 transcripts, DNMT3B expression was significantly downregulated in androgenetic and parthenogenetic haploid groups when compared the biparental control embryos (Figure 8A), suggesting that developmental events that require *de novo* methylation may be impaired in haploid embryos. In contrast, unaltered expression levels of DNMT1 and TET1 indicate that DNA methylation maintenance and active demethylation is not affected in both androgenetic and parthenogenetic haploid groups.

**Figure 8.**
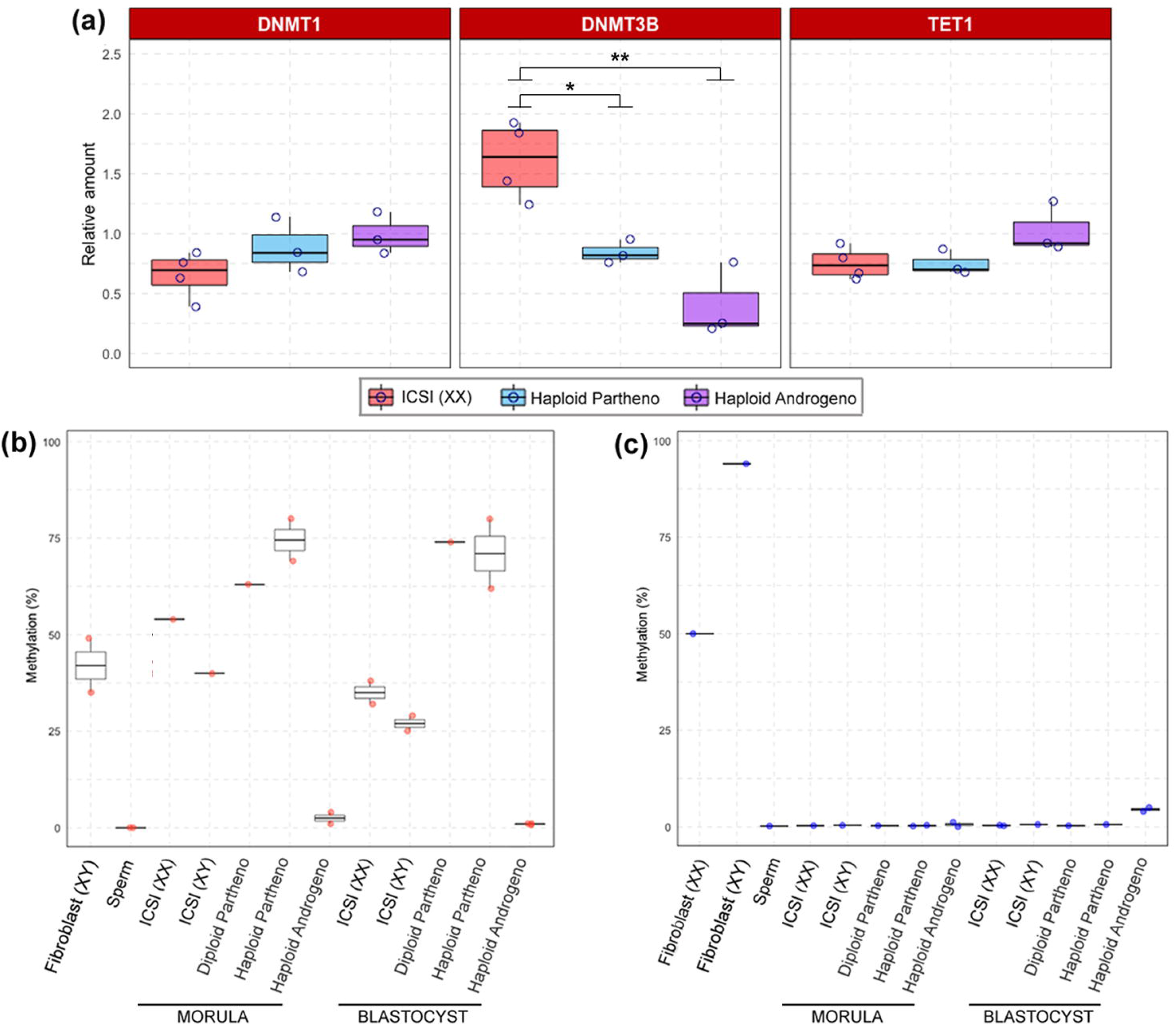
Relative gene expression of DNA methylation-related enzymes and DNA methylation profiles of XIST and KCNQ1 DMR. (a) Relative gene expression of DNMT1, DNMT3B and TET1 in biparental diploid female (ICSI), haploid parthenogenetic and androgenetic embryos. ICSI (XX), intracytoplasmic sperm injection using female sorted-sperm; ICSI (XY), intracytoplasmic sperm injection using male sorted-sperm; Haploid Partheno: haploid parthenogenetic embryo obtained by oocyte activation using ionomycin followed by cyclohexymide; Haploid Androgeno, haploid androgentic embryo obtained by ICSI + oocyte enucleation; Fibroblast (XX), female fibroblast cells; Fibroblast (XY), male fibroblast cells.

### Methylation patterns of the XIST and KCNQ1OT1 DMRs are unaltered in hAE

Finally, to evaluate if the abnormal expression of X chromosome and KCNQ1 locus genes was associated to alterations in methylation patterns, we performed bisulfite sequencing of the XIST and KCNQ1 DMRs in morula and blastocyst stage embryos. As observed in other mammalian species, the bovine XIST DMR region was 50% methylated in adult female fibroblasts and hypermethylated in adult male fibroblasts, supporting the notion of dosage-compensation by X chromosome inactivation in female somatic tissues (Figure 8B). On the other hand, the XIST DMR was hypomethylated in sperm, male and female embryos. Besides, since the XIST DMRs in haploid androgenotes, and parthenogenetic (haploid and diploid) embryos were all hypomethylated, these results indicate that the upregulation of XIST expression in hAE is not related to the methylation status of its DMR.

As expected for DMRs controlling imprinted loci, the KCNQ1 DMR was hypomethylated in the male gamete and approximately 50% methylated in fibroblast and in male and female biparental (ICSI-derived) morula and blastocyst stage embryos (Figure 8C). Diploid and haploid parthenogenetic embryos at the morula and blastocysts stage showed an elevated methylation levels of the KCNQ1 DMR, indicating a hypermethylation of the maternal allele. In contrast, androgenetic embryos showed hypomethylated pattern at morula and blastocyst stages that resembled the patterns observed in spermatozoa that are typical of this imprinted locus. These results suggest that the abnormal expression patterns of genes from the KCNQ1 locus in hAE, i.e. KCNQ1OT1 and PHADL2, may at least in part be due to an epigenetic dysregulation resulting from the exclusive presence of the paternal allele.

## DISCUSION

Here, we analyzed different methodologies for production of bovine hAE and the potential causes for their limited ability to develop to the blastocyst stage. Our results indicate anomalies in gene expression of X chromosome and of the KCNQ1 imprinted loci, suggesting the involvement of epigenetic regulators in the developmental constraints of haploid androgenotes in the bovine species.

Initially, we evaluated the removal of the oocyte spindle (enucleation) before and after in vitro fertilization on the embryonic development (Latham et al., 2002; Lagutina et al., 2004; Vichera et al., 2011). The embryos produced by these methods showed similar cleavage rates, but blastocyst development was seriously limited. In agreement with ours results, previous studies have reported that most of the bovine hAE are arrested after first cell divisions and that only a few of them can reach the blastocyst stage (Lagutina et al., 2004; Vichera et al., 2011). In mice, hAE produced by fertilization of enucleated oocytes also showed limited blastocyst development (11%) compared to the IVF group (90%) (Kono et al., 1993). Nonetheless, our results showed that their developmental potential was enhanced when the enucleation was performed post-IVF. In agreement with these findings others have indicated that enucleation during the telophase stage of second meiosis (TII) shows advantages over enucleation at the metaphase II stage. For instance, TII enucleation allows the removal of smaller ooplasm fragments (Bordignon and Smith, 1998; Lee and Campbell, 2006), can be performed in the absence of UV irradiation (Kuznyetsov et al., 2007; Sagi et al., 2019), and it also allows the selection of the best oocytes for enucleation through the exclusive use oocytes that respond promptly to fertilization by second polar body extrusion (Kuznyetsov et al., 2007).

However, although the production of hAE was feasible by using conventional in vitro fertilization (IVF), the analysis of pronuclear formation showed a high proportion of multinucleated zygotes, indicative of polyspermic fertilization, which has been previously reported in mice (Kono et al., 1993) and cattle (Lagutina et al., 2004). Polyspermic fertilization after bovine IVF can vary between 5% to 25% (Roh et al., 2002; Coy et al., 2005), which makes it an unreliable tecnique for producing haploid embryos. To assure the effective monospermic fertilization we used ICSI. In agreement with previous reports (Latham et al., 2002; Lagutina et al., 2004; Vichera et al., 2011; Yang et al., 2012), our results showed that haploid androgenetic zygotes can be reliably produced by combining ICSI and oocyte enucleation.

Cleavage rate and cell number of hAE after 48 h of culture indicated that micromanipulation (ICSI and enucleation) does not influence initial mitotic divisions of early embryonic development. Nonetheless, although androgenetic haploidy does not impact development up to around the 3^rd^ - 4^th^ mitotic division, the hAE underwent developmental arrest zygotic genome activation (ZGA) at the 8-cell stage. In addition, androgenetic embryos that progressed beyond ZGA underwent a second arrest at the morula stage. Similar results have been reported in cattle (Winger et al., 1997; Vichera et al., 2011), sheep (Matsukawa et al., 2007), mouse (Kono et al., 1993; Latham et al., 2002; Hu et al., 2015a; Hu et al., 2015b), and human species (Kuznyetsov et al., 2007; Sagi et al., 2019). Further development to the blastocyst stage, of both haploid parthenotes and androgenotes was severely limited compared to diploid embryos, evidencing a deleterious effect of haploidy on the very early stage of embryogenesis.

Since reports in mice have indicated that the presence Y- and/or the absence of a X-chromosome in haploid androgenetic embryos restricts development beyond the four-cell stage (Latham et al., 2002; Yang et al., 2012), we used sex-sorted sperm to produce bovine haploid androgenotes. When using Y-chromosome sorted sperm, development was arrested at a very early stage before compaction, confirming previously murine studies comparing X- and Y-chromosome carrying androgenotes. Similarly, we showed that bovine haploid androgenotes derived from X-chromosome sorted sperm develop poorly and only rarely reach the blastocyst stage. Moreover, since sex sorting techniques are typically 90% accurate (Sharpe and Evans, 2009), these results indicate that the poor development of haploid androgenotes produced using X-chromosome sorted sperm cannot be explained by erroneous use of Y- chromosome sperm. Moreover, assessment of total cell number and blastocyst morphology in haploid androgenotes, which are positively correlated with blastocyst quality (Sagirkaya et al., 2006; Kong et al., 2016), showed fewer cells and delayed blastulation compared not only to diploid controls but also to the haploid parthenotes, suggesting that the haploid androgenetic condition is less capable to support early development than haploid gynogenetic condition. On the other hand, the analysis of chromosomal constitution showed that aneuploidy levels were higher in haploid parthenotes than in androgenotes, excluding chromosomal segregation errors as a cause of the limited embryonic development in the haploid androgenotes. Since bovine centrosomes are inherited from the sperm at fertilization and are responsible for organizing the mitotic spindle of the zygote (Long et al., 1993; Navara et al., 1994; Navara et al., 1995; Navara et al., 1996; Sutovsky et al., 1996a; Sutovsky et al., 1996b), it is likely that because the sperm centrosome is the only responsible for spindle formation after the removal of the oocyte’s spindle the hAE maintain a stable karyotype during the subsequent mitotic divisions.

According to Matsukawa *et al.* (2007), hAE that undergo early arrest commonly present micronuclei and picnotic nuclear formation. Because of this, we wanted to evaluate the nuclear morphology of early arrested zygotes. The hAE, ICSI and parthenogenetic-derived embryos showed similar rates of anomalies, suggesting that micromanipulation procedures (ICSI/enucleation and chemical oocyte activation) are related to the early development defects. In agreement with these results, numerous studies have associated bovine ICSI with several developmental anomalies such as insufficient sperm head decondensation (Rho et al., 1998; Malcuit et al., 2006), delayed pronuclear formation (Aguila et al., 2017), and altered gene expression (Arias et al., 2015). However, although such ICSI harmful effects certainly can contribute to the early developmental failure of androgenotes, they do not explain the development arrest the occurs beyond ZGA. Accordingly, a study by Latham et al. (2002) indicated that the injection procedure (ICSI) does not influence in vitro development and excluded chromosomal abnormalities as the main cause for the limited development of haploid androgenetic mouse embryos. Altogether, the results above support the notion that the reduced developmental potential of bovine haploid androgenotes is likely related to gene expression anomalies occurring after ZGA.

The effects of paternal haploidy on expression of imprinted genes during early embryogenesis has not yet been evaluated in the bovine species. Uniparental haploid embryos possess one copy of either the paternal or maternal genomes, thus theoretically, the expression of imprinted genes would be either present or undetectable relative to biparental embryos. The silencing of one of the X chromosomes in diploid female embryos is regulated by the expression of Xist, a non-coding RNA that acts as a major effector on X-chromosome inactivation (XCI). The methylation of Xist prevents its expression and many have studied the potentially negative effects on development of parthenogenetic (Chen et al., 2019) and cloned (Zeng et al., 2016) mammalian embryos. On the other hand, the KCNQ1 imprinted domain is one of the largest known imprinted clusters, and its altered imprinting has been associated with fetal overgrow or large offspring syndrome (LOS) (Lee et al., 1999; Chen et al., 2015). This region is regulated by the KvDMR1 located in the promoter of the non-coding KCNQ1OT1 gene which is maternally methylated. Kcnq1ot1 is paternally expressed and negatively regulate the expression of several maternally expressed genes, including CDKN1C, KCNQ1, and PHLDA2 (Ager et al., 2008). Thus, differential expression of X-linked and imprinted genes, can help to address the causes behind the limited in vitro developmental potential of haploid androgenotes. Our data revealed similar expression levels at the time of ZGA among groups, where only CDKN1C was upregulated in androgenotes compared to the other groups, indicating a differential alteration at time of ZGA in imprinted gene expression. At the morula stage, XIST was highly expressed in the hAE and the X-linked genes PGK1 and HPRT showed similar levels among groups. In cattle, the presence of XIST transcripts has been reported as early as the 2-cell stage (Mendonca et al., 2019). Microarray and RNA-seq analyses of bovine blastocysts demonstrated higher expression of X-linked genes in female compared with male embryos, indicating that dosage compensation initiates later (Bermejo-Alvarez et al., 2010; Min et al., 2017). A recent report has indicated that XIST accumulation and XCI in bovine embryos starts at the morula stage. However, XIST colocalization with repressive marks (H3 lysine 27 trimethylation) on histones was only detected by day 7 blastocysts, indicating that complete XCI is only partially achieved at the blastocyst stage (Yu et al., 2020). Although *XIST* accumulation did not lead to globally reduced expression of X-linked genes, and X-Chr inactivation is only partially achieved at the blastocyst stage (Bermejo-Alvarez et al., 2010; Yu et al., 2020), the effects of the dysregulated expression of this long-noncoding RNA in haploid androgenetic morulas and blastocyst stages needs further investigation to clarify its impacts on chromosome-wide downregulation of gene expression. Latham et al. (2002) reported elevated expression of Xist RNA in haploid mouse androgenotes, and a similar pattern for the PGK1 gene, suggesting that haploid androgenotes may undergo deficient XCI, or that the embryos that initiate the XCI process begin to die soon thereafter. Haploid androgenotes with the greatest degree of Xist RNA expression, PGK1 gene repression, and repression of other X-linked genes may die within a narrow period of time just after ZGA, which is consistent with our findings that the majority of haploid androgenotes fail to progress to the morula and blastocyst stage.

To our knowledge, this is the first study analyzing the expression of genes that belonging to the KCNQ1 imprinted domain in haploid androgenetic mammalian embryos. By analogy with X inactivation in the mouse species, the KCNQ1OT1 is paternally expressed as early as two-cell stage and maintained throughout preimplantation development, but the ubiquitously imprinted genes KCNQ1 and CDKN1C are paternally repressed at the morula/blastocyst stage. By contrast, placentally imprinted genes TSSC4 and CD81 show biallelic expression in the blastocyst (Umlauf et al., 2004; Lewis et al., 2006). In this study, the KCNQ1OT1 and PHLDA1 were overexpressed in haploid androgenetic morula stage embryos. Also, CDKN1C expression was unexpectedly upregulated in androgenotes compared to male diploid embryos. As discussed earlier, haploid androgenotes that were able to progress up to morula stage might have escaped from KCNQ1OT1-silencing or those with the greatest degree of imprinting repression arrested just after ZGA. Thus, the abnormal expression of the KCNQ1 imprinted domain could potentially affect the development and differentiation of haploid androgenetic early stage embryos. Finally, IGF2R and GNAS, two maternal imprinted genes (Jiang et al., 2015), showed similar expression levels, which suggests that their imprinting was relaxed, or as previously reported, the monoallelic expression in ruminants may not be required for most imprinted genes during early embryonic development (Cruz et al., 2008).

DNA cytosine methylation is one of the most important modifications in the epigenetic genome and plays essential roles in various cellular processes, including genomic imprinting, X chromosome inactivation, retrotransposon silencing, as well as regulation of gene expression and embryogenesis (Reik et al., 2001). The addition of methyl groups to cytosine residues is catalyzed by DNA methyltransferases (DNMT1 for maintenance and DNMT3A and DNMT3B for de novo methylation (Pablo J. Ross, 2018). Active DNA demethylation has been ascribed to TET activity (Iqbal et al., 2011). In cattle, the presence of DNMT3B has been reported as the major responsible for the control of methylation levels at advanced preimplantatory stages (Pablo J. Ross, 2018). Besides, among the factors required for demethylation process, TET1 is the predominant expressed enzyme after zygote genome activation (Bakhtari and Ross, 2014). Our results indicate that haploid and biparental embryos had similar levels of DNMT1 and TET1 transcripts. The zygotically expressed form of DNMT1 maintains the methylation of imprints at each cell cycle during early embryonic development (Hirasawa et al., 2008; Kurihara et al., 2008), suggesting that hAE are able to maintain their methylation imprints. Conversely, DNMT3B was deficient in both haploid groups. In accordance with our results, a previous study in mice demonstrated that the production and derivation of androgenetic haploid ESCs were severely impaired when Dnmt3b was deficient (He et al., 2018), suggesting that proper Dnmt3b activity and the content of methylation is essential for the development of mammalian haploid androgenetic embryos. Moreover, embryogenesis is severely impaired when Dnmt3b homozygous deletion (Okano et al., 1999). In mouse embryos, the inactivation of Dnmt3b induces a partial global hypomethylation, and even though the catalytic activities of DNMT3b and DNMT3a can compensate for each other, DNMT3B makes a greater contribution to the methylome, specifically in a set of CpG-dense sequences associated with pluripotency and developmental imprinted genes (Kato et al., 2007; Auclair et al., 2014). In addition, DNMT3B also has specific roles in the methylation of many CpG islands on autosomes and the inactive X chromosome that are dramatically hypomethylated in Dnmt3b KO embryo (Auclair et al., 2014). Nonetheless, it remains unknown whether bovine haploid androgenetic embryos undergo global hypomethylation.

Finally, we performed a gene specific bisulfite sequencing in order to analyze whether DMR methylation patterns were related to the high expression of XIST and KCNQ1OT1. As for the XIST DMR, all groups were demethylated, which is in accordance with its biallelic expression at during the early stages of bovine embryogenesis (Bermejo-Alvarez et al., 2010; Yu et al., 2020). In bovine sperm cells, the *XIST* gene does not appear methylated (Mendonca et al., 2019), suggesting that *XIST* would be expressed in androgenetic cells It is likely that the XIST DMR undergoes methylation later during embryonic development, as the onset of XCI initiates at blastocyst stage (Yu et al., 2020). As for the KCNQ1 DMR, embryos carrying a maternal allele (biparental and parthenogenetic embryos) were beyond 60% methylated. However, since embryos from the haploid androgenetic group were demethylated and resembled the imprinted profile observed in the sperm (Robbins et al., 2012), if such hypomethylation is associated or it is in part responsible for the altered gene expression of the KNCQ1OT1 gene, needs to be further investigated.

In conclusion, this study has shown that micromanipulation effects and chromosomal abnormalities are not main factors affecting the development of bovine hAE. On the other hand, we show that the failure of haploid androgenetic bovine embryos to develop to the blastocyst stage is associated with abnormal expression of key factors involved in DNA methylation, XCI and genomic imprinting, suggesting that their early developmental constraint is regulated at an epigenetic level. In order to obtain a better understanding of epigenetic regulation in the mammalian haploid androgenetic model, future studies will be aimed at a more in-depth analysis of the global epigenetic features. This will involve investigation by global transcriptomic and methylation analysis as well the study of repressive epigenetics marks in haploid embryos.

## Author Contributions

LA, JT, and LS contributed to conception and design of the study. JS, MG and AG contributed with experimental procedures. LA, JT and LS wrote the manuscript. All authors contributed to manuscript revision, read, and approved the submitted version.

---

Supplemental figure 1

Proportion of cleaved embryos at the 8-cell stage at 48h of culture. IVF, in vitro fertilized; ICSI, intracytoplasmic sperm injection using female-sorted semen; Haploid partheno, haploid parthenogenetic embryos obtained by oocyte activation using ionomycin followed by cyclohexymide; Haploid androgeno, haploid androgenetic embryo obtained by ICSI + oocyte enucleation.

Supplemental figure 2

Morphological assessment of haploid androgenetic embryos produced with sperm carrying Y-chromosome at 144 h of culture. Representative (a) morphologies and (b, c) nuclear counts of embryos harvested at Day-6 (144 h) of culture.

