## Supplementary figures and images for "Dysregulated gene expression of imprinted and X-linked genes: a link to poor development of bovine haploid androgenetic embryos"

### Supplemnetal Figure 1

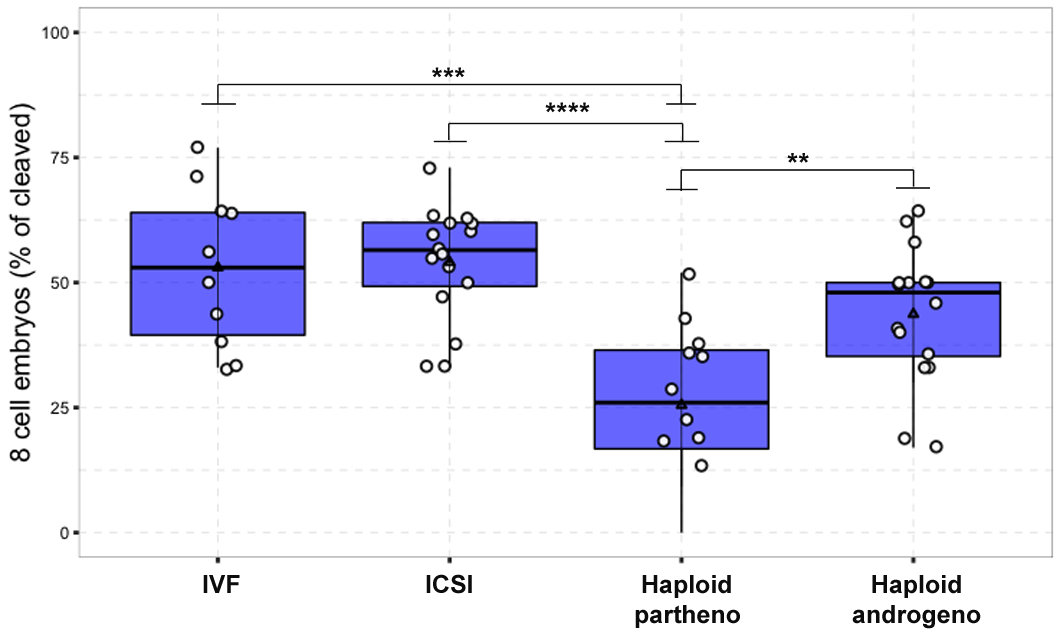

### Supplemnetal Figure 2

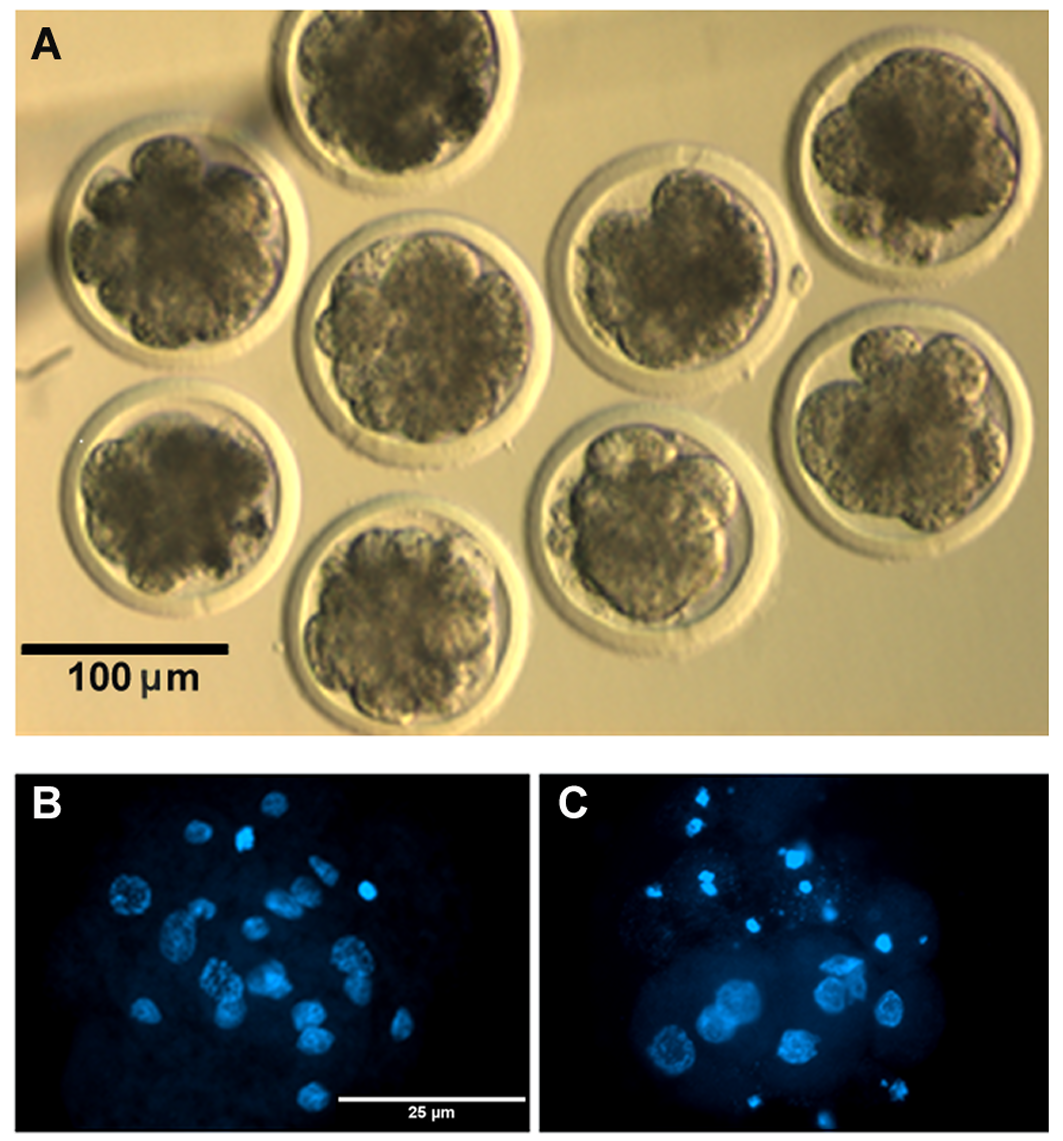
